# Sex and APOE Genotype Differentially Shape Microglial Transcriptomic Profiles Across the Hippocampus and Cortex

**DOI:** 10.64898/2026.08.31.748388

**Authors:** Sandra Ledesma-Corvi, Samantha A Blankers, Andrew J McGovern, Morgan Towriss, Jennifer Kim, Annie V Ciernia, Liisa AM Galea

## Abstract

Female sex and the APOEε4 allele are top risk factors for Alzheimer’s disease (AD). Microglia play a role in the pathogenesis of AD, yet how sex and APOE genotype affect microglia remain poorly understood. Here, we characterized the transcriptomic and morphological profiles in the hippocampus and cortex of microglia from humanized APOEε3 and APOEε4 mice of males and females. The hAPOEε4 genotype was associated with sex-dependent effects on microglia co-expression modules in both brain regions, involving cellular stress and immunometabolism processes. In females, a hippocampal cell cycle module was supported by reduced microglial proliferation. Cortical modules were enriched for lipid metabolism and immune-related processes whose expression decreased in females but increased in male hAPOEε4. Male, but not female, hAPOEε4 microglia shifted toward an ameboid state in both regions. Together, these findings reveal sex-dependent microglial responses to APOEε4 across brain regions and highlight the need to incorporate sex-specific approaches into AD research.

## INTRODUCTION

Alzheimer’s disease (AD) is a progressive neurodegenerative disorder characterized by cognitive decline, with hallmark pathological features including amyloid beta (A*β*) plaque accumulation and neurofibrillary tau tangles, accounting for 60-70% cases of dementia worldwide.^1^ In addition to advancing age, the greatest non-modifiable risk factors for sporadic AD are female sex and Apolipoprotein E (APOE) ε4 genotype.^2,3^ Although longevity plays a role in sex differences in lifetime risk for AD, human females also exhibit higher levels of AD neuropathology and faster cognitive decline compared to human males with AD.^4–6^ In addition, APOE accounts for approximately 25% of the total heritability of sporadic AD and represents the strongest genetic risk factor for late-onset AD, with 3-15 fold increased risk with one to two copies of the APOEε4 allele.^7^ APOEε4 genotype is also associated with increased amyloid deposition, neurofibrillary tangle load and dysfunction of the medial temporal lobe, which is one of the first regions to display atrophy with AD.^8–10^ Notably, the impact of APOEε4 genotype is stronger in females, as female APOEε4 carriers exhibit an increased risk of earlier AD onset and greater AD neuropathology, including phosphorylated tau accumulation and more severe cognitive impairment compared to male APOEε4 carriers.^2,11,12^ Collectively, these observations suggest that female sex and APOE genotype independently and interactively affect AD risk and endophenotypes, highlighting the importance of understanding the mechanisms underlying these interactions to advance precision medicine in AD.

Neuroinflammation plays a fundamental role in the progression of AD neuropathology.^13^ Genome-wide association studies reveal that many genetic risk factors for late-onset AD are exclusively or highly expressed in microglia in the central nervous system (CNS),^14^ highlighting microglia as central players in the neuroinflammatory cascade of AD pathology. Microglia, which are the resident immune cells within the CNS, actively surveil the brain parenchyma and respond to neurodegeneration through the recognition of misfolded proteins, phagocytosis of debris, and the orchestration of immune response via chemokine/cytokine release.^15^ Microglia play a protective role early in the response to AD-related amyloidosis,^15,16^ however, in later AD stages chronic activation of microglia may be detrimental, perpetuating a cycle of neuronal damage and degeneration.^17^ Intriguingly, in human females but not males, microglia activation mediates the progression of AD pathogenesis,^18^ suggesting it is essential to consider sex effects on microglia along with APOE genotype to understand AD progression.

Sex differences in microglial transcriptional signatures have also been reported in mouse models of AD, with female microglia progressing faster over activated response microglia (ARM) trajectory^19^ and displaying greater disease associated microglia (DAM) phenotype, including sex-specific enhancement of interferon signaling in response to Aβ.^20^ Female microglia also exhibit different metabolic profiles, being more glycolytic and less phagocytic than male microglia.^21,22^ APOE genotype is an important modulator of microglia activation, with the APOEε4 allele linked with increased microgliosis and neuroinflammation,^23^ impaired phagocytosis, and accumulation of lipid droplets (LD) driving a maladaptive and damaging LD-accumulating microglia (LDAM) in response to Aβ.^24,25^ Although some studies have shown that APOE genotype and sex interact to affect microglial transcriptional profiles^26^ and plaque coverage^27^ in animal models of AD; studies investigating their interactive effects on microglia remain limited, particularly at early stages of pathology.

The objective of this study was to characterize sex differences in the influence of APOEε4 genotype on microglial phenotype in two key AD-vulnerable regions, the hippocampus (HC) and cortex (CTX). We hypothesized that APOEε4 genotype would differentially impact microglial phenotype in a sex-dependent manner, displaying different transcriptomic signatures and morphological alterations in each brain region. To test this hypothesis, microglia were isolated from the HC and CTX of male and female seven month old humanised (h) APOEε3 and hAPOEε4 mice.^28^ Weighted gene co-expression network analysis (WGCNA) and differentially expressed gene (DEG) analysis were used to identify sex-dependent effects of APOE genotype on microglial transcriptomic signatures in each region, followed by pathway enrichment analysis. Microglial proliferation was quantified and microglial morphological states were assessed to examine potential differences in microglial activation state in both regions.

## RESULTS

### Identification of co-expression modules showing a sex by genotype interaction in hippocampal microglia

We performed WGCNA to identify microglial transcriptomic signatures associated with APOE genotype in a sex-dependent manner in the HC (Fig 1A). The analysis used the 50% most variable genes (7,593 genes) detected in hippocampal-isolated microglia. Then, we explored the relationship between module eigengenes and experimental group traits (Fε4, Fε3, Mε4 and Mε3). We identified 16 co-expression modules, of which 8 showed significant correlations with group traits (Fig 1B, 1C). Given our interest in exploring sex-dependent microglial signatures associated with hAPOEε4 in the HC, we next evaluated the effects of sex, genotype and their interaction on each module eigengene (ME) using a linear model. Modules showing a significant sex by genotype interaction were selected for downstream analysis, including functional annotation, identification of the top 10 intramodular hub genes, and microglia gene list enrichment analysis to assess enrichment for previously described microglial gene signatures.^29^ We identified four modules (magenta, black, grey60 and greenyellow) showing a significant sex by genotype interaction, that were selected for further characterization (see Table S2). The greenyellow module was not significantly enriched for any GO terms or pathways and was not investigated further. The remaining three modules were enriched for specific pathways and functions as described in detail below.

**Figure 1.**
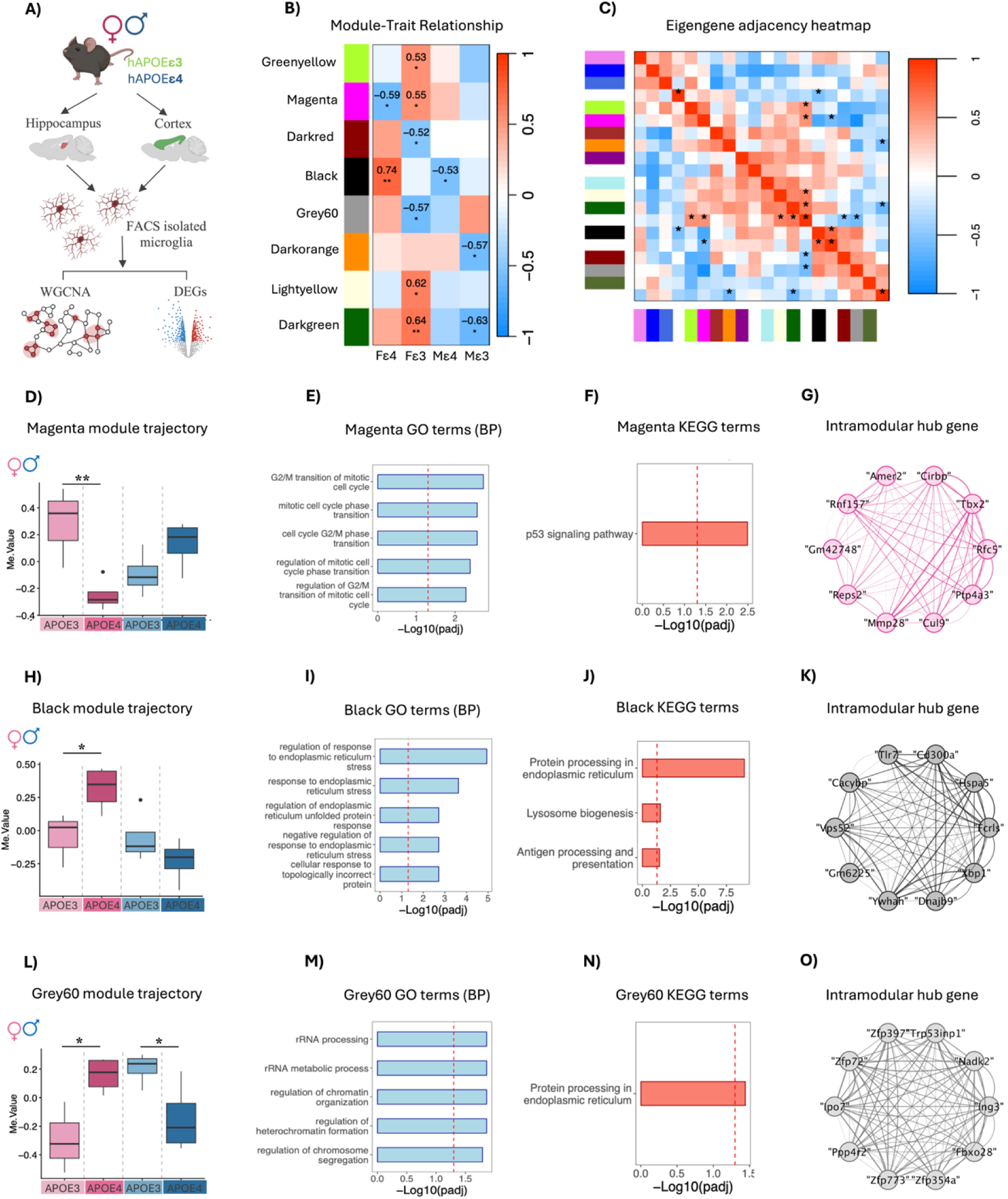
Identification of sex-dependent gene modules associated with hAPOEε4 in hippocampal microglia. **(A)** Experimental design for the analysis of FACS-sorted microglia from the hippocampus and cortex of 7-month old male and female hAPOEε3 and hAPOEε4 mice. **(B)** The correlations between module eigengene (ME) and the traits of interest: female hAPOEε4 (Fε4), female hAPOEε3 (Fε3), male hAPOEε4 (Mε4) and male hAPOEε3 (Fε3). The values in the heatmap are Pearson’s correlation coefficients and stars represent significant correlations: *p < 0.05; ** p < 0.01. Positive correlations are shown in red, whereas negative correlations are shown in blue; the intensity of the color corresponds to the strength of the correlation. **(C)** Heatmap of the eigengene adjacency matrix. Each row and column in the heatmap corresponds to one module eigengene (labeled by color) or a trait of interest. Within the heatmap red color represents positive correlation, while blue color represents negative correlation. **(D, H, L)** Module eigengene boxplot in the **(D)** magenta, **(H)** black and **(L)** grey60 modules across APOE genotype and sex. Results of Šídák’s multiple comparisons posthoc analyses are denoted. \**p* < 0.05 and \*\**p* < 0.01. **(E, I, M)** The top 5 Gene Ontology (GO) enriched Biological Processes (BP) in the **(E)** magenta, **(I)** black and **(M)** grey60 module. The dotted line indicates the threshold for BH-corrected *p* < 0.05. **(F, J, N)** The top 5 Kyoto Encyclopedia of Genes and Genomes (KEGG) enriched terms in **(F)** magenta, **(J)** black and **(N)** grey60 module. The dotted line indicates the threshold for BH-corrected *p* < 0.05. **(G, K, O)** Network plot with the top 10 intramodular hub genes in the **(G)** magenta, **(K)** black and **(O)** grey60 module. See also Figure S2 and Table S2.

### The cell cycle-associated magenta module is reduced in female hAPOEε4 and was enriched for biological processes related to the cell cycle

For the magenta module, there was a significant sex by genotype interaction (F(1,11) = 14,451, *p* = 0.003). Šídák-corrected posthoc analysis revealed that the ME of the magenta module was significantly decreased in hAPOEε4 females compared with hAPOEε3 females (estimate = -0.535, *p* = 0.007), while no significant effects were observed in males (*p* = 0.24; Fig 1D and Table S2). Genes in this module were enriched in biological processes related to the cell cycle, including G2/M phase transition and mitotic phase transition (Fig 1E, BH-corrected *p* < 0.05). Cellular component analysis also showed enrichment in nuclear membrane, focal adhesion, and outer kinetochore (Table S2, BH-corrected *p* < 0.05). The p53 signaling was the only enriched KEGG pathway identified, and top hub genes included *Cirbp*, *Tbx2, Cul9*, all of which are involved in cell cycle phase transition and mitotic nuclear division^30,31^ (Fig 1F, 1G). Consistently, MGEnrichment analysis revealed enrichment of the magenta module for previously described microglial signatures, including Ki67+ > Ki67-microglia signatures among others (Fig S2A, FDR-corrected *p* < 0.05). Together, these results suggest a coordinated expression of genes involved in cell cycle regulation, potentially reflecting reduced microglial proliferation in hAPOEε4 females compared to hAPOEε3 females.

### hAPOEε4 genotype reduces microglial proliferation in the dorsal HC of female mice

To validate the cell cycle-related pathways identified in the magenta module, BrdU+/Iba1+ co-expressing cells were quantified to examine potential differences in microglial proliferation in the dorsal and ventral HC of male and female hAPOEε4 and hAPOEε3 mice (Fig 2). Although the repeated-measures ANOVA revealed only marginal effects of sex by region (dorsal vs ventral) by genotype (*p* = 0.094), based on the transcriptomic findings showing a decrease of cell cycle-related module in hAPOEε4 females, we hypothesized *a priori* that female hAPOEε4 mice would have lower levels of microglial proliferation in the HC. Consistent with this hypothesis, hAPOEε4 females showed a lower density of BrdU+/Iba1+ cells compared with hAPOEε3 females in the dorsal HC (*a priori*: *p* = 0.0145, Cohen’s *d* = 1.139; Fig 2A), but not in the ventral HC (*p* = 0.257; Fig 2B). No significant effects were observed in males across regions (*p’s >* 0.294).

**Figure 2.**
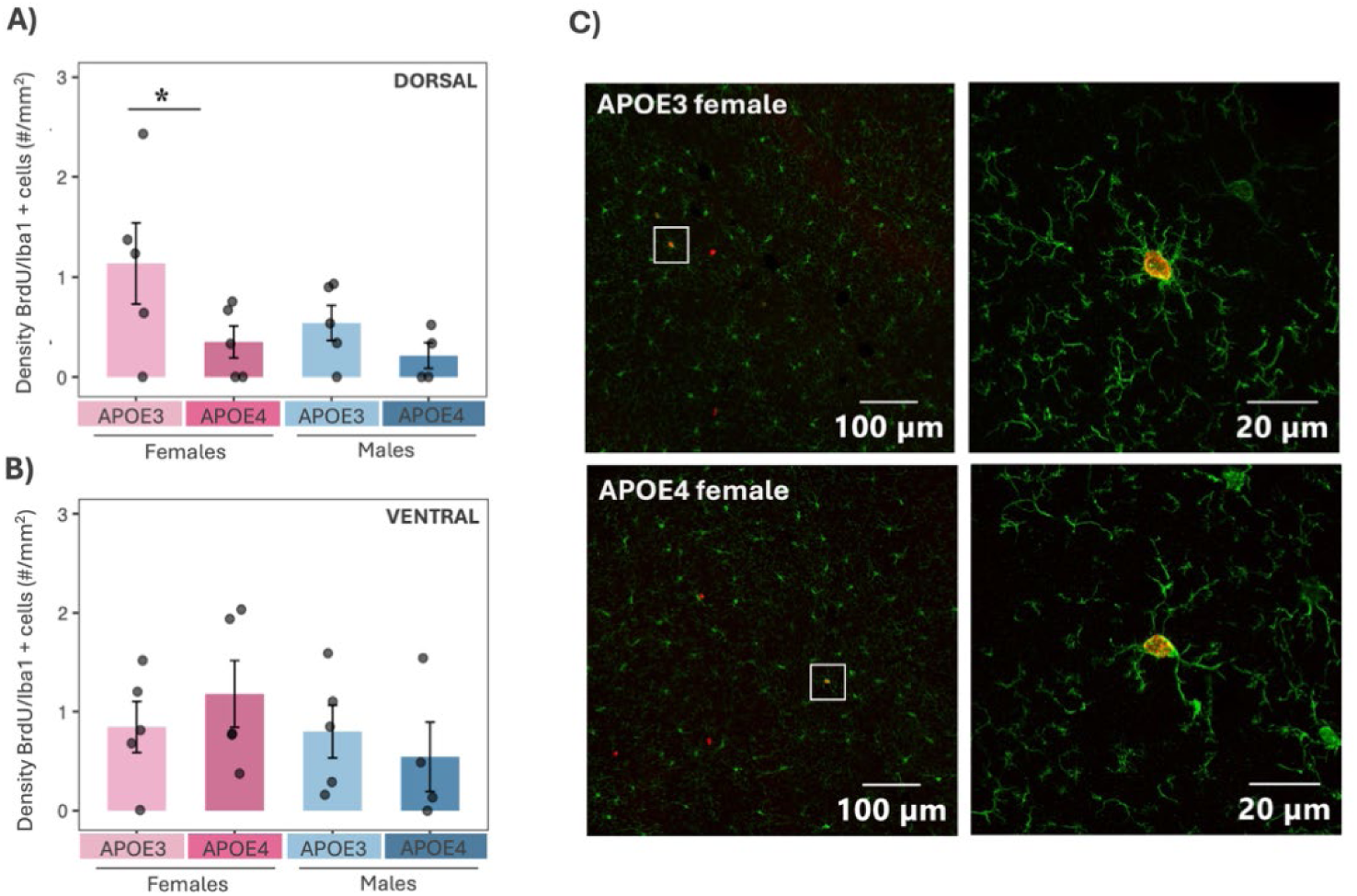
Microglia proliferation decreases in the hippocampus of hAPOEε4 females (A,B) Average density of BrdU/Iba1+ cells in the **(A)** dorsal and **(B)** ventral hippocampus. Data represent average mean ± standard error of the mean (SEM). Individual symbols are shown for each animal. **(C)** Representative image showing the colocalization of Iba1+ (green) and BrdU+ (red) cells in the dorsal hippocampus of female hAPOEε3 and hAPOEε4 mice. Scale bars are 100 μm and 20 μm.

### Modules enriched in endoplasmic reticulum stress response and rRNA metabolic process were increased in female, but not male, hAPOEε4 mice

We also identified the black module showing a significant sex by genotype interaction (F(1,11) = 8.062, *p* = 0.016). Šídák-corrected posthoc analysis showed that black ME was significantly increased in hAPOEε4 females compared with hAPOEε3 females (estimate = 0.366, *p* = 0.047), but not in males (*p* = 0.369) (Fig 1H and Table S2). Genes in this module were enriched for biological processes related to the regulation of endoplasmic reticulum (ER) stress, protein folding, and macroautophagy (Fig 1I, BH-corrected *p* < 0.05). Cellular component analysis showed enrichment in the ER chaperone complex, lysosomal lumen and membrane, as well as phagocytic vesicles, while the molecular function analysis revealed enrichment in protein-folding, chaperone binding, and ubiquitin protein ligase binding among others (Table S2, BH-corrected *p* < 0.05). KEGG pathways were also related with protein processing in ER, lysosome biogenesis, and antigen processing and presentation (Fig 1J, BH-corrected *p* < 0.05). The top hub genes in this module, included *Hspa5*, *Dnajb9* and *Xbp1* (Fig 1K), are also involved in the ER stress response and unfolded protein response (UPR).^32–34^ Consistently, MGEnrichment analysis revealed enrichment of the black module for previously described microglial signatures, including microglial cellular stress and microglial sensome signatures (Fig S2A, FDR-corrected *p* < 0.05). Together, these results suggest an enhanced coordinated expression of genes managing ER proteostasis and cellular stress in the microglia of hAPOEε4 females.

Finally, we also identified the grey60 module showing a sex by genotype interaction (F(1,11) = 17.139, *p* = 0.002). Šídák posthoc test revealed that grey60 ME was significantly increased in hAPOEε4 females compared with hAPOEε3 females (estimate = 0.452, *p* = 0.018) and decreased in hAPOEε4 males compared with hAPOEε3 males (estimate = -0.354, *p* = 0.043) (Fig 1L and Table S2). Genes in this module were enriched in biological processes related to rRNA metabolic process and regulation of heterochromatin formation (Fig 1M, BH-corrected *p* < 0.05).

Consistently, cellular component analysis showed enrichment in the ISWI-type complex and ribosomes, while the molecular function revealed enrichment in histone binding, ribosome binding and p53 binding (Table S2, BH-corrected *p* < 0.05). The protein processing in ER was the only enriched KEGG pathway identified, and top hub genes included *trp53inp1*, which encodes an antiproliferative and proapoptotic protein involved in cell stress response and acts as a dual regulator of transcription and autophagy (Fig 1N, 1O, BH-corrected *p* < 0.05).

### Identification of sex-dependent hAPOEε4-related differentially expressed genes in hippocampal microglia

We next performed a differentially expressed gene (DEG) analysis using a generalized linear model including sex, genotype and their interaction. In the HC, we identified 6 significantly decreased and 2 significantly increased genes in hAPOEε4 females compared to hAPOEε3 females (q-value < 0.05; Fig 3A); and 10 significantly decreased and 4 significantly increased genes in hAPOEε4 males compared to hAPOEε3 males (q-value < 0.05; Fig 3B). Finally, we identified four genes *Nfil3*, *Cribp*, *Epha2* and *Hsph1* showing a significant sex by genotype interaction (q-value <0.05; Fig 3C).

**Figure 3.**
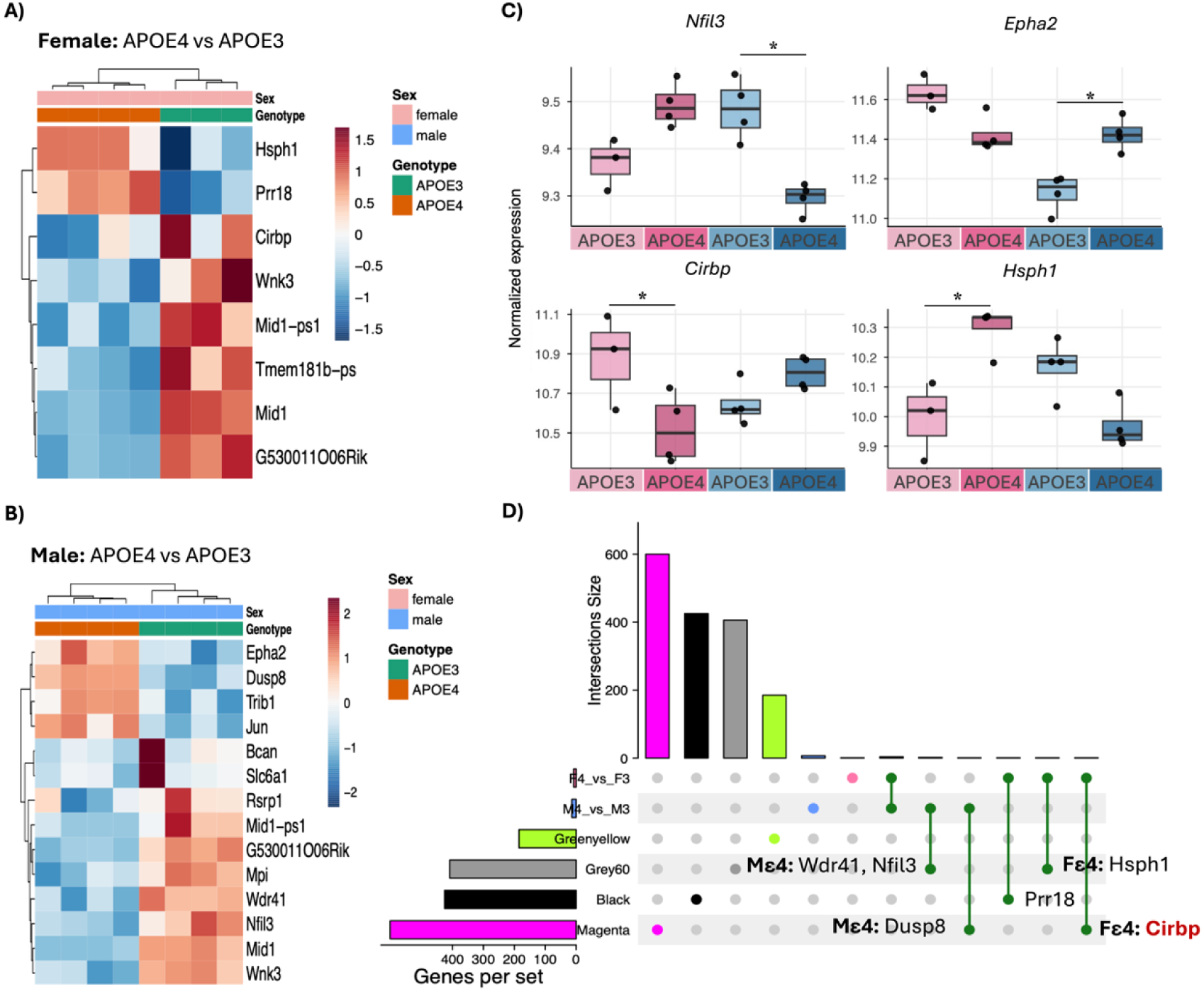
Differential Gene Expression analysis and overlap with sex-dependent hAPOEε4-associated modules in hippocampal microglia. **(A,B)** Heatmap of normalized gene expression values (z-score, VST) for genes differentially expressed by APOE genotype (APOE4 vs. APOE3) within **(A)** females and **(B)** males (q-value < 0.05) in hippocampal microglia. Each row is a gene, and each column is a sample. Samples hierarchically cluster by APOE genotype. **(C)** Boxplots of normalized gene expression values (VST) for genes showing a significant sex by genotype interaction (interaction term, q-value < 0.05). Individual symbols are shown for each animal. Simple comparison of APOE genotype (APOE4 vs. APOE3) within each sex are denoted: * q-value < 0.05. **(D)** UpSet plot illustrating the overlap between DEGs identified and previously selected WGCNA gene modules in the hippocampus. Gene names are shown for overlapping genes, with red highlighting genes identified as intramodular hub genes within the corresponding module. See also Table S3.

We next examined whether any of the DEGs overlapped with the WGCNA modules showing a significant sex by genotype interaction. Of the 22 DEGs identified, 6 genes were distributed across significant modules, with three located in the grey60 module, one in the black module and two in the magenta module (Fig 3D). Within the grey60 module we identified *Wdr4*1 and *Nfil3* genes which were significantly decreased in hAPOEε4 males compared to hAPOEε3 males (*Wdr41*: logFC = -0.395; q-value = 0.02; *Nfil3*: logFC = -0.948; q-value = 0.044), as well as the *Hsph1* gene which was significantly increased in hAPOEε4 females compared with hAPOEε3 females (logFC = 0.651; q-value = 0.033). The gene *Prr18,* which was increased in hAPOEε4 females compared with hAPOEε3 females (logFC = 1.18; q-value = 0.006), was identified within the black module, which was associated with biological processes related to endoplasmic reticulum stress and protein folding.

Finally, *Cirbp* and *Dusp8* were identified within the magenta module. Notably, *Cirbp* was identified as a hub gene within this module and was significantly decreased in hAPOEε4 females compared with hAPOEε3 females (logFC = -0.605; q-value = 0.042). In contrast, *Dusp8* was increased in hAPOEε4 males compared to hAPOEε3 males (logFC = 1.04; q-value = 0.042). Consistent with the functional enrichment of the magenta module, both genes have been associated with pathways involved in cell cycle regulation.^31,35^

### Identification of co-expression modules showing a sex by genotype interaction in cortical microglia

We next performed a WGCNA to identify microglial transcriptomic signatures associated with APOE genotype in a sex-dependent manner in the CTX. The analysis used the 50% most variable genes (7,520 genes) detected in cortical-isolated microglia. Using the same approach described in the HC, we identified 32 co-expression modules, of which 18 showed significant correlations with traits of interest (Fε4, Fε3, Mε4 and Mε3; *p* < 0.05) (Fig 4A, 4B). Seven modules (red, brown, lightsteelblue1, green, cyan, darkmagenta and orangered4) showed significant sex by genotype interactions (see Table S4) and were selected for downstream analysis, including functional annotation, the identification of the intramodular hub genes, and microglia gene list enrichment analysis. The orangered4 and green modules were not significantly enriched for any GO term or pathways and were not investigated further. The remaining five modules were enriched for specific pathways and functions as described in detail below.

**Figure 4.**
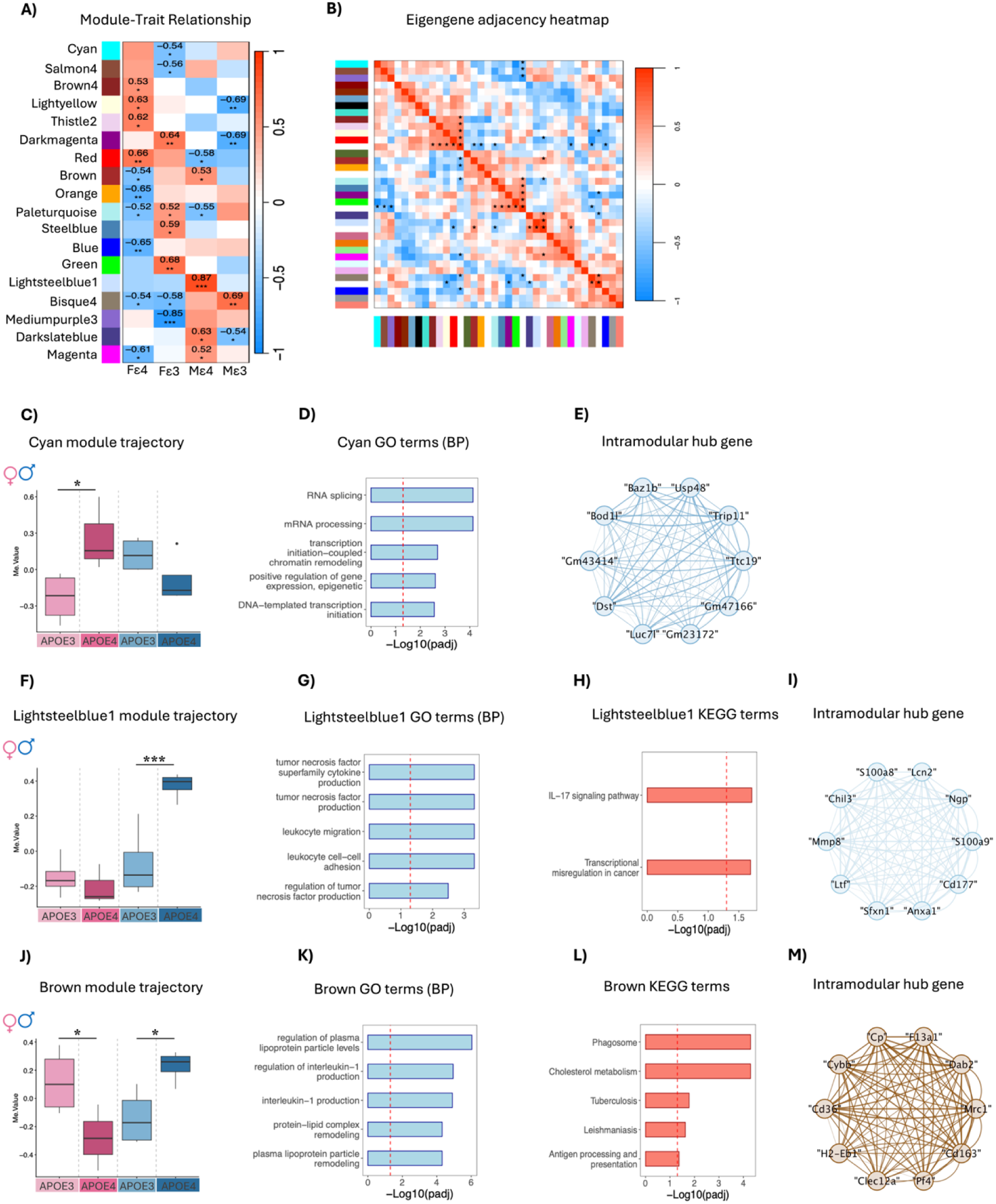
Identification of sex-dependent gene modules associated with hAPOEε4 in cortical microglia. **(A)** The correlations between module eigengene (ME) and the traits of interest: female hAPOEε4 (Fε4), female hAPOEε3 (Fε3), male hAPOEε4 (Mε4) and male hAPOEε3 (Fε3). The values in the heatmap are Pearson’s correlation coefficients and stars represent significant correlations: *p < 0.05; ** *p* < 0.01; *** *p* < 0.001. Positive correlations are shown in red, whereas negative correlations are shown in blue; the intensity of the color corresponds to the strength of the correlation. **(B)** Heatmap of the eigengene adjacency matrix. Each row and column in the heatmap corresponds to one module eigengene (labeled by color) or a trait of interest. Within the heatmap red color represents positive correlation, while blue color represents negative correlation. **(C, F, J)** Module eigengene boxplot in the **(C)** cyan, **(F)** lightsteelblue1 and **(J)** brown modules across APOE genotype and sex. Results of Šídák’s multiple comparisons posthoc analyses are denoted. \**p* < 0.05 and \*\*\**p* < 0.001. **(D, G, K)** The top 5 Gene Ontology (GO) enriched Biological Processes (BP) in **(D)** cyan, **(G)** lightsteelblue1 and **(K)** brown modules. The dotted line indicates the threshold for BH-corrected *p* < 0.05. **(H, L)** The top 5 Kyoto Encyclopedia of Genes and Genomes (KEGG) enriched terms in **(H)** lightsteelblue1 and **(L)** brown modules. The dotted line indicates the threshold for BH-corrected *p* < 0.05. **(E, I, M)** Network plot with the top 10 intramodular hub genes in the **(E)** cyan, **(I)** lightsteelblue1 and **(M)** brown modules. See also Figure S2-3 and Table S4.

### Female hAPOEε4 is associated with increased expression of mRNA metabolism-enriched cyan module, with a similar trend in interferon-β-related module

We identified the cyan module showing a significant sex by genotype interaction (F(1,11) = 10.05, *p* = 0.009). Šídák posthoc test revealed that the ME was significantly increased in hAPOEε4 females compared with hAPOEε3 females (estimate = 0.489, *p* = 0.023), but not in males (*p* = 0.342) (Fig 4C and Table S4). Functional annotation revealed enrichment in biological processes related to RNA splicing and mRNA metabolism (Fig 4D, BH-corrected *p* < 0.05), whereas no significant KEGG pathways or MGEnrichment were observed. Among the top hub genes in the module, we also identified *Luc7l* (Fig 4E), which is involved in RNA splicing,^36^ overall suggesting an enhanced coordinate gene expression involved in mRNA processing and splicing in microglia from hAPOEε4 females.

The red module also showed a significant sex by genotype interaction (F(1,11) = 5.21, *p* = 0.043). Šídák posthoc analysis showed a trend toward increased red ME in hAPOEε4 females compared with hAPOEε3 females (estimate = 0.136, *p* = 0.06) (Fig S2A and Table S4), but not in males (*p* = 703). Genes in this module were enriched for biological processes related to response to interferon-beta (Fig S3B, BH-corrected *p* < 0.05), while no significant KEGG pathways or MGEnrichment were detected. Notably, among the top hub genes in the module we identified *Ddx3x* (Fig S3C), which has been previously linked to interferon-beta signaling.^37^ Interestingly, among the hub genes, we also identified three X chromosomally encoded transcripts (*Kdm5c*, *Eif2s3x* and *Ddx3x),* all of which have previously been associated with X-linked intellectual disability^38,39^ (Fig S3C).

### Male hAPOEε4 is associated with increased expression of neuroimmune-**related LightSteelBlue1 module**

The lightsteelblue1 module showed a significant sex by genotype interaction (F(1,11) = 12.598, *p* = 0.004). Šídák posthoc analysis showed that the ME was significantly higher in hAPOEε4 males compared with hAPOEε3 males (estimate = 0.447, *p* = 0.001), but not in females (*p* = 0.834) (Fig 4F and Table S4). Functional annotation revealed enrichment in biological processes related to neuroinflammatory and immune response (Fig 4G, BH-corrected *p* < 0.0 5), consistent with cellular compartment enrichment in membrane raft and specific granule (Table S4, BH-corrected *p* < 0.05). Furthermore, KEGG annotation revealed that genes in this module play an important role in the IL-17 signaling pathway (Fig 4H, BH-corrected *p* < 0.05). Among the top hub genes in the module, we identified *S100A8*, *S100A9, Lcn2* and *Anxa1*(Fig 4I), which play prominent roles in the regulation of inflammatory processes and immune response.^40–42^ Consistently, MGEnrichment analysis revealed enrichment of the lightsteelblue1 module for genes increased in P14 Hdac1/2-deficient microglia compared with WT microglia (Fig S2B, FDR-corrected *p* < 0.05).

We also identified the darkmagenta module showing a significant sex by genotype interaction (F(1,11) = 7.894, *p* = 0.017). However, no significant Šídák posthoc comparisons were detected, although a trend was observed in males (estimate = 0.284, *p* = 0.08; females: *p* = 0.215) (Fig S3D and Table S4). Genes in this module were enriched for biological processes related to T cell differentiation (Fig S3E, BH-corrected *p* < 0.05), whereas no significant KEGG pathways or MGenrichments were observed.

### Female APOEε4 decreased, whereas male APOEε4 increased, expression in a module enriched in lipid metabolism and neuroinflammation

Finally, the brown module showed a significant sex by genotype interaction (F(1,11) = 13.871, *p* = 0.003). Particularly, Šídák posthoc analysis revealed that brown ME was significantly decreased in hAPOEε4 females compared with hAPOEε3 females (estimate = -0.397, *p* = 0.045) and increased in hAPOEε4 males compared with hAPOEε3 males (estimate = 0.366, *p* = 0.046) (Fig 4J and Table S4). Genes in this module were enriched for biological processes related to lipid metabolism and neuroinflammation (Fig 4K, BH-corrected *p* < 0.05), consistent with cellular compartment enrichment in lipid transport structures, the endosomal-lysosomal system, and immune-related structures. At the molecular function level, enrichment terms included lipoprotein and lipid binding, cholesterol and sterol binding, immune receptor activity (e.g., pattern recognition and scavenger receptors), and amyloid-beta binding (Table S4, Bh-corrected *p* < 0.05). KEGG pathways were also related with phagosome, cholesterol metabolism, and antigen processing and presentation (Fig 4L, BH-corrected *p* < 0.05). Consistent with the module’s functional annotation, the top hub genes included *Cd36* (Fig 4M), a scavenger receptor that has been reported to mediate microglial response to β-amyloid,^43^ as well as cholesterol and fatty acid metabolism.^44^ Notably, *Apoe*, *Lpl* and *clec7a* previously identified as DAM markers,^45^ although not identified as hub genes, were also present within this module, overall suggesting a sex-dependent coordinated expression of genes involved in microglia lipid metabolism and immune response in hAPOEε4 mice.

Consistently, MGEnrichment analysis revealed enrichment of the brown module for previously described microglial signatures, including neurodegenerative microglia signatures, among others (Fig S2B, FDR-corrected *p* < 0.05).

### Identification of sex-dependent hAPOEε4-related differentially expressed genes in cortical microglia

In the CTX, we identified 8 significantly decreased and 1 significantly increased genes in hAPOEε4 females compared to hAPOEε3 females (q-value < 0.05 Fig 5A) and 7 significantly decreased and 3 significantly increased genes in hAPOEε4 males compared to hAPOEε3 males (q-value < 0.05; Fig 5B). Finally, we identified six genes with significant sex by genotype interaction, including *Clec7a*, *Zfp142* and *Fzd7* (q-values < 0.05; Fig 5C).

**Figure 5.**
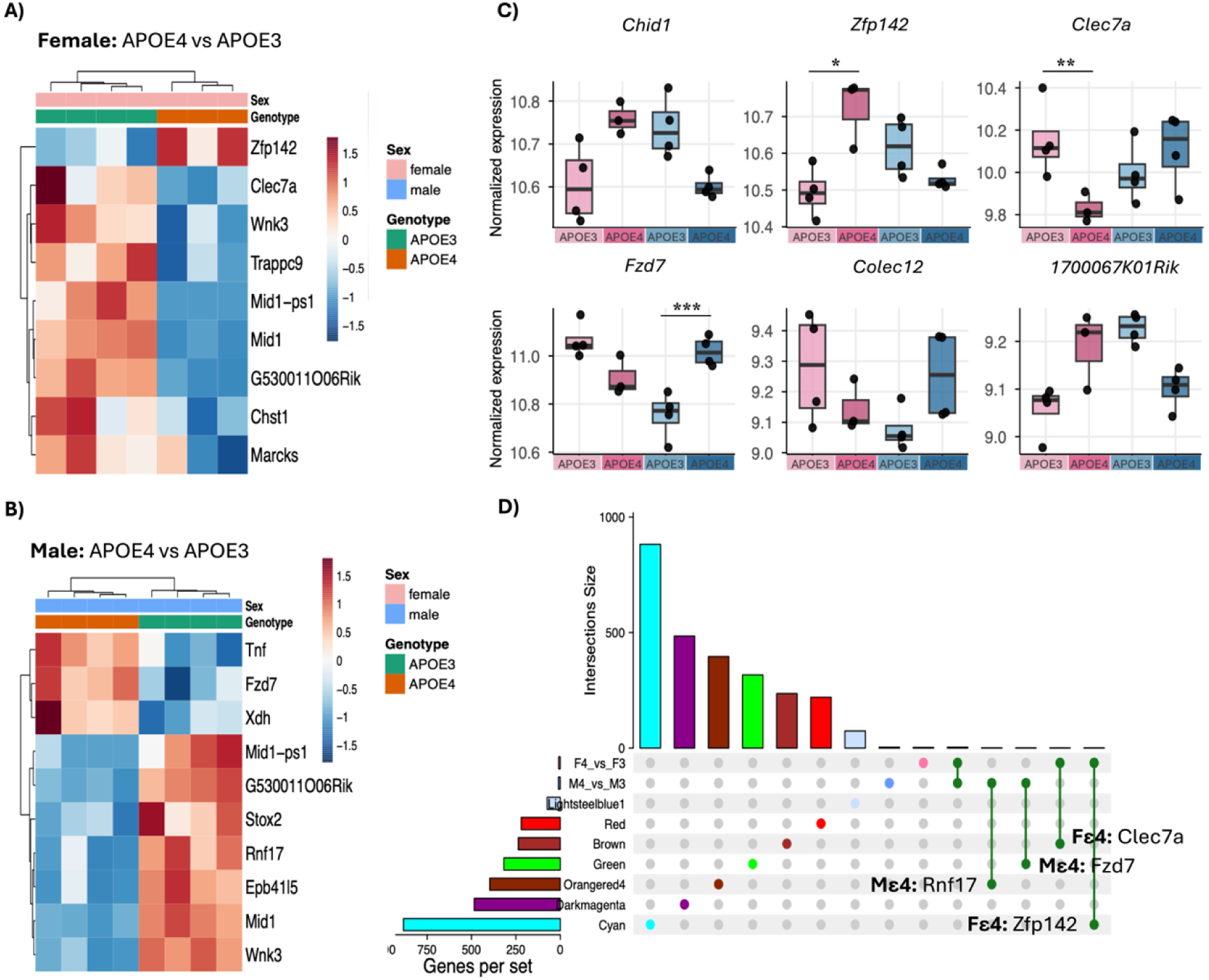
Differential Gene Expression analysis and overlap with sex-dependent hAPOEε4-associated modules in cortical microglia. (A,B) Heatmap of normalized gene expression values (z-score, VST) for genes differentially expressed by APOE genotype (APOE4 vs. APOE3) within **(A)** females and **(B)** males (q-value < 0.05) in cortical microglia. Each row is a gene, and each column is a sample. Samples hierarchically cluster by APOE genotype. **(C)** Boxplots of normalized gene expression values (VST) for genes showing a significant sex by genotype interaction (interaction term, q-value < 0.05). Individual symbols are shown for each animal. Simple comparison of APOE genotype (APOE4 vs. APOE3) within each sex are denoted: * q-value < 0.05, ** q-value < 0.01 and *** q-value < 0.001. **(D)** UpSet plot illustrating the overlap between DEGs identified and previously selected WGCNA gene modules in the cortex. Gene names are shown for overlapping genes. See also Table S5.

Next, we explored the overlap between DEGs and WGCNA modules showing a significant sex by genotype interaction in the CTX. Of the 19 DEGs identified, four were distributed across significant modules (Fig 5D), including one in the brown module, one in the green, one in the orangered4 and one in the cyan module.

Notably, the *Clec7a* gene, that was decreased in hAPOEε4 females compared with hAPOEε3 females (logFC = -0.853; q-value = 0.001), was identified within the brown module. Consistent with our findings in WGCNA, *Clec7a* is a pattern recognizing receptor which has previously been identified as a key marker of DAM.^45^ Within the cyan module we identified the *Zfp142* gene, a transcriptional regulator, which was significantly increased in hAPOEε4 females compared with hAPOEε3 females (logFC = 0.366; q-value = 0.035). The *Rnf17* gene, which was significantly decreased in hAPOEε4 males compared with hAPOEε3 males (logFC = -1.868; q-value < 0.001), was identified within the orangered4 module. Finally, within the green module we identified the *Fzd7* gene, which was significantly increased in hAPOEε4 males compared with hAPOEε3 males (logFC = 0.41; q-value < 0.001).

### Sex-dependent microglia morphological alterations associated with hAPOEε4 across brain regions

Given that transcriptomic analyses showed sex-dependent hAPOEε4 transcriptional signatures enriched in neuroinflammation, immune response, lipid metabolism and ER stress-related pathways, and given that microglia morphology is often associated with their functional activity, we further examined whether these transcriptomic signatures were accompanied by changes in microglial morphology. Using MicrogliaMorphologyR,^46^ we quantified shifts in morphological populations within the same brain regions assessed for microglia RNAseq from female and male hAPOEε3 and hAPOEε4 mice. Using unbiased clustering, we identified four clusters of microglia morphology in both regions: ramified, ameboid, hypertrophic, and rod-like (Fig 6A and Fig S4). In the HC, there was a significant interaction between APOE genotype, sex and cluster in the ventral (*X*^2^(3, *n* = 20) = 19.011, *p* < 0.001; Fig 6B) but not dorsal (*p* = 0.07) region. Post-hoc analyses revealed that hAPOEε4 males, but not females, showed an increase in ameboid microglia (estimate = 0.294, Šídák-corrected *p* = 0.0012) and a decrease in ramified microglia (estimate = -0.216, Šídák-corrected *p* = 0.025) compared with hAPOEε3 male mice, showing a shift from ramified to ameboid microglia state with hAPOEε4 in males (Fig 6B). In the CTX, we also observed a significant interaction between APOE genotype, sex and cluster (*X*^2^(3, *n* = 20) = 13.321, *p* = 0.004). Similarly to HC, post-hoc analyses revealed that hAPOEε4 males showed more ameboid microglia (estimate = 0.208, Šídák-corrected *p*= 0.007) and fewer ramified microglia (estimate = -0.185, Šídák-corrected *p* = 0.01) than males hAPOEε3 (Fig 6C), showing a shift from ramified to ameboid microglia state with hAPOEε4 in males. The percentage of hypertrophic and rod-like microglia remained consistent across APOE genotypes and sexes in both brain regions.

**Figure 6.**
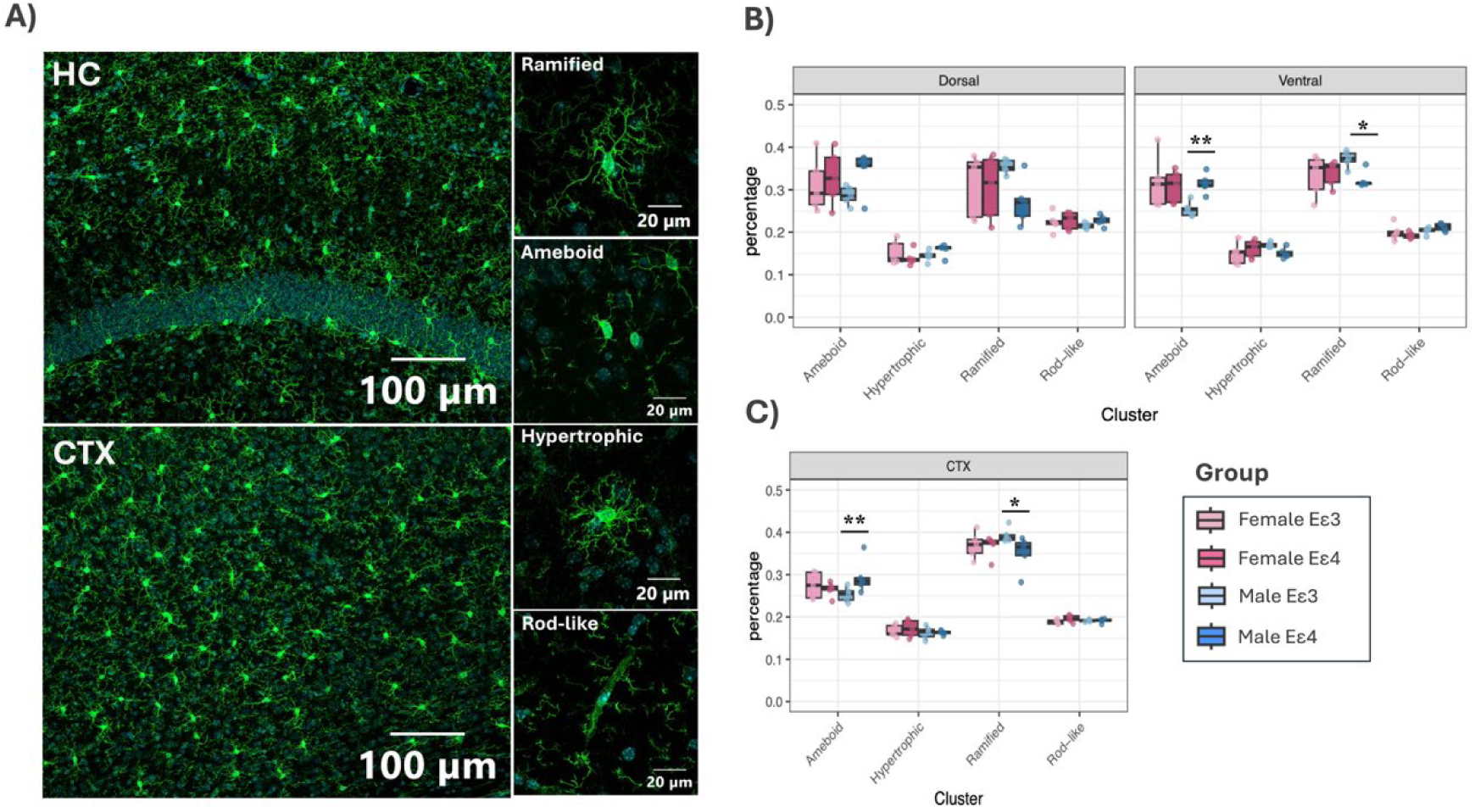
Microglia morphology shifts toward an ameboid state in hAPOEε4 males. **(A)** Example of immunofluorescent images of hippocampal and cortical microglia cells and the different morphological state identified, stained with Iba1 marker (Green). Nuclei are stained with DAPI (blue). Scale bars are 100 μm and 20 μm. **(B,C)** Percentage of microglia within each morphological cluster between genotype and sex in **(B)** the dorsal and ventral hippocampus and **(C)** cortex. Individual symbols are shown for each animal. Results of Šídák’s multiple comparisons posthoc analyses are denoted. \**p* < 0.05; ** *p* < 0.01. See also Figure S4 and Table S6.

## DISCUSSION

Using an unbiased approach, the present study demonstrates striking sex-specific microglial signatures in the hippocampus and cortex of adult mice that are dependent on APOE genotype. Here, we show that hAPOEε4 genotype is associated with transcriptional alterations that differ between sexes in microglia in both the HC and CTX, with females and males exhibiting distinct signatures compared to their respective hAPOEε3 controls at 7 months of age. Consistent with the transcriptome findings, hAPOEε4 genotype altered microglial proliferation and morphology in a sex-specific manner, with females, but not males, showing reduced microglial proliferation in the HC compared with hAPOEε3 female mice.

Furthermore, the morphology of microglia shifted towards a more ameboid and less ramified microglial state in both brain regions in hAPOEε4 compared to hAPOEε3 males, but not in females. Together, these findings highlight sex differences in microglial transcriptomic and morphological signatures across brain regions in a model of AD risk at an early time point, underscoring the need to incorporate sex-specific approaches within AD research and treatment development.

### hAPOEε4 genotype-associated increases in an ER stress-related module in hippocampal microglia in females but not in males

In the HC, hAPOEε4 females exhibited altered microglial signatures characterized by an increase in an ER stress response-related module eigengene, characterized by hub stress-related genes *Hspa5* and *Xbp1,* and enriched for stress-related microglia gene signatures. Together, these findings suggest ER stress-related transcriptional alterations in female microglia with the hAPOEε4 genotype. This is consistent with previous studies showing that APOEε4 induces ER stress in microglia,^47^ increases eIF2α phosphorylation, a key downstream effector of PERK-mediated ER stress signaling,^48^ and augments a microglial subset characterized by cellular stress.^49,50^ Although transient activation of the unfolded protein response can restore microglia homeostasis, chronic ER stress can impair cellular function and viability, promote inflammatory responses, and contribute to disease progression.^51^ Therefore, future studies are needed to determine whether these early transcriptomic changes represent a transient, adaptive response or contribute to progressive microglia dysfunction in hAPOEε4 females.

### hAPOEε4 genotype-associated decreases in a cell cycle-related module and reduced microglia proliferation in females but not in males

Female hAPOEε4 microglia showed reduced expression of a cell cycle-related module eigengene, with *Cirbp* identified as the hub gene. Notably, *Cirbp* is a stress-responsive gene that positively modulates cell cycle progression and proliferation by accelerating G0/G1 and G1/S phase transition through the regulation of different factors such as HuR and various cyclins.^31,35^ Consistent with this transcriptional profile, microglia proliferation was reduced in hAPOEε4 females, whereas no significant effects were observed in males. The functional implications of this early sex-specific reduction in microglia proliferation remain unclear. Enhanced proliferation represents a key feature of the early microglial response to acute injuries and chronic neurodegeneration.^52,53^ Thus, decreased microglial proliferation could compromise microglia functions during disease progression. Conversely, reduced microglia proliferation could represent an adaptive response to prevent dysfunctional or stressed microglia from proliferating. Previous studies in males have shown that proliferating microglia exhibit transcriptional alterations in a mouse model of familial AD and reduced capacity for amyloid-beta clearance at 6 months of age.^54^ Moreover, reducing microglia proliferation in male mice improves cognitive decline and prevents synaptic degeneration in AD models.^52,55^ However, it is important to consider that the majority of these studies were conducted in male mice or without sex stratification and in transgenic animal models involving various AD gene mutations. Therefore, further studies are needed to better understand the functional implications of this early reduction in microglia proliferation observed in hAPOEε4 females at early stages of the disease.

### hAPOEε4 genotype-associated sex-dependent alterations of a lipid metabolism- and immune-related module in cortical microglia

In the CTX, a distinct pattern of microglia transcriptional changes was observed, with sex differences emerging in microglial signatures characterized by an altered module eigengene levels related to lipid metabolism and immune function. This module was decreased in hAPOEε4 females and increased in hAPOEε4 males.

This is consistent with previous studies showing distinct microglia-specific endophenotypes between female and male AD subjects, with females showing a decrease in lipid metabolism pathways.^56^ Interestingly, this module was also significantly enriched in microglia neurodegenerative (MGnD) signatures,^57,58^ a microglia phenotype that arises in response to degenerating/apoptotic neurons and is regulated by the TREM2-APOE pathway.^57^ Consistent with this enrichment, *Clec7a*, a key marker of DAM/MGnD,^45^ was significantly decreased in hAPOEε4 females. The MGnD phenotype has been proposed to limit pathology by clearing cellular debris and other threats,^57^ but little is known about sex differences in MGnD microglia in the context of AD risk. Here, our results in females align with human studies showing suppression of MGnD genes in the brain of female AD donors carrying the APOE4 allele compared to individuals homozygous for the APOE3 allele.^59^ In addition, previous studies in mice have shown a negative role of microglia APOE4 in the induction of the MGnD response to neurodegeneration at 8 month of age, and that deletion of APOE4 restores the MGnD phenotype associated with neuroprotection in P301S tau transgenic mice and decreases pathology in APP/PS1 mice.^59^ However, it is also important to mention that other studies have reported a different pattern, with female microglia displaying greater DAM/MGnD phenotypes than male microglia in both AD models^20^ and aged mice.^60,61^ These differences could be due to the transgenic animal model used (e.g., 5xFAD) or the age of the mice, since here we focus on characterizing the effects of hAPOEε4, the greatest genetic risk factor for late-onset sporadic AD, at an earlier time point.

### hAPOEε4 genotype-associated increases in an inflammatory- and immune-related module in cortical microglia in males but not in females

In the CTX, microglia from hAPOEε4 males also showed an increase in a neuroinflammatory and immune-related module eigengene, including IL-17 signaling and tumor necrosis factor (TNF) superfamily cytokine production, with inflammatory-related hub genes *S100A8* and *S100A9.*^42^ In line with these results, previous studies have shown elevated IL-17 signaling in microglia from male AD subjects compared to controls,^56^ and that APOEε4 increases TNF-α secretion in primary microglia.^48^Together these results suggest a sex-dependent effect of hAPOEε4 genotype on inflammatory and immune-related signatures in cortical microglia, with a more prominent effect in males.

### hAPOEε4 genotype alters microglia morphology in a sex-dependent manner across both brain regions

Given that microglia function is typically associated with morphological changes, we next quantified microglia morphological differences across both brain regions. Here, we found that the hAPOEε4 genotype induced a morphological shift toward a more ameboid and less ramified microglial state in males, but not in females, across both brain regions. This is consistent with previous studies showing that APOEε4 microglia have smaller overall cell size and shorter average branch length compared to APOEε3 microglia,^48^ and with observations in animal models of AD where microglia exhibit an ameboid morphology,^21,62^ a state typically associated with increased reactivity in response to damage or pathological stimuli in the brain.^63^ Previous studies from humans also found that microglia from male AD patients showed more ameboid appearance with process retraction, while cells from females AD patients were more complex and variable in morphology.^21^ Ameboid cells are associated with more phagocytic capacity^64^ and elevated secretion of pro-inflammatory cytokines such as TNF-α,^65^ which are in line with our transcriptomic results in the cortex, suggesting a pro-inflammatory microglia state in hAPOEε4 males. Interestingly, despite the transcriptomic changes observed in hAPOEε4 females, these were not accompanied by detectable changes in microglial morphology in either brain region. This dissociation is consistent with recent evidence showing that DAM microglia can display both ameboid and ramified morphology, indicating that morphology and transcriptional state are not always tightly coupled.^66^ Together these findings suggest that microglia morphology and transcriptional signatures may not always align in capturing microglia alterations, particularly in females.

### Limitations of the study

Several limitations of this study should be noted. First, our study is restricted to seven months of age, a time point prior to middle age. It is increasingly recognized that microglia are highly heterogeneous cells that play influential roles across the lifespan, which may cumulatively affect AD risk. Therefore, studies at later time points may reveal divergent or convergent microglial responses, and further studies will be needed to assess whether these sex differences resolve or progress with age. In this respect, microglia show changes in humans across aging with shifts to more monocyte infiltration after middle age, suggesting different patterns will emerge later in life with one study showing some subtle variations by sex.^67,68^ Second, we used hAPOEε4 mice, which represent a genetic risk model for AD rather than a model of overt AD pathology. However, this model provides a valuable opportunity to examine sex differences in the effects of APOEε4 alone on microglial phenotypes, given that APOEε4 is the strongest genetic risk factor for late-onset AD^7^ and the impact of APOEε4 genotype is greater in females.^2,11,12^ Third, we focused on transcriptomic and morphological microglial signatures in hAPOEε4 mice, but we did not evaluate cognitive performance or neuropathological outcomes. Therefore, whether the sex-dependent microglial alterations observed in this model are associated with or contribute to AD-related pathology remains to be determined.

### Conclusions

Together, these findings showed that hAPOEε4 genotype affected the microglial transcriptomic signatures and morphological states in a sex-dependent manner across brain regions prior to middle age. Future work should expand our knowledge of how sex and hAPOEε4 genotype affect microglia function and brain health by exploring how these early microglia signatures may influence AD endophenotypes, as well as examining later time points such as middle age. Overall, our findings underscore the need to incorporate and analyse with sex as a biological variable in AD research. Given that hAPOEε4 alleles and female sex are among the top non-modifiable risk factors for AD, understanding how the interplay between these two factors contribute to AD risk and affect microglia function will be critical to informing targeted and effective treatments.

## RESOURCE AVAILABILITY

### Lead contact

Further information and requests for resources and reagents should be directed to and will be fulfilled by the lead contact, Liisa Galea and Annie Ciernia.

### Material availability

This study did not generate new unique reagents.

### Data and code availability

Raw files and count matrix are available on NCBI GEO at GSE312921 This paper does not report original code

Any additional information required to reanalyze the data reported in this paper is available from the lead contact upon request.

## ACKNOWLEDGEMENTS

We are deeply grateful to Kimberly Go and Stephanie Lieblich for their technical assistance. This work was supported by Cure of AD (to L.A.M.G) and CIHR Postdoctoral Fellowship (to S.L.C PTA-208351).

## AUTHORS CONTRIBUTIONS

Conceptualization, S.L.C., L.A.M.G. and A.V.C.; methodology, S.L.C., S.B., L.A.M.G. and A.V.C.; investigation, S.L.C., S.B., M.T., J.K. and A.J.M.; formal analysis, S.L.C. L.A.M.G. and A.V.C.; resources, L.A.M.G. and A.V.C.; funding acquisition, L.A.M.G. and A.V.C.; writing - original draft, S.L.C., S.B., M.T., L.A.M.G. and A.V.C.; supervision, L.A.M.G. and A.V.C. All authors read and approved the final manuscript.

## DECLARATION OF INTEREST

## METHODS

### EXPERIMENTAL MODEL AND STUDY PARTICIPANT DETAILS

#### Mice

Humanized (h)APOEε3 and (h)APOEε4 mice were obtained from Taconic Bioscience via Cure for Alzheimer’s Fund. These lines were generated by targeted replacement of murine *APOE* gene with human *APOEε3* and *APOEε4* alleles in embryonic stem cells followed by blastocyst injection.^28^ Resulting chimeras were then backcrossed with C57BL/6 mice for seven generations and the resulting mice were housed and bred at the University of British Columbia (UBC). Mice were housed in plastic cages (36.5 x 20.5 x 14 cm) in groups of 5-9 on a 12:12 hour light/dark cycle and provided food and water ad libitum until they reached 7 months of age. Male and female *APOE* mice at 7 months of age were used for all studies. For transcriptomic studies (n=4 mice/genotype/sex), mice were anesthetized with isoflurane and transcardially perfused with 1X HBSS and inhibitor cocktail (5 μg/mL Actinomycin D, 10 μM Triptolide, and 27.1 μg/mL Anisomycin). Brains were extracted, and the CTX and HC were dissected separately on ice and stored temporarily in transport buffer (1XHBSS + inhibitor cocktail) until microglia isolation. For morphology studies (n=5 mice/genotype/sex), mice were anesthetized with isoflurane and transcardially perfused with 30 mL of 0.9% saline and 30 mL of freshly made 4% paraformaldehyde. Brains were then extracted, post-fixed in 4% paraformaldehyde for 24h, transferred to 30% sucrose for cryoprotection, and stored at 4°C until tissue slicing. For validation studies (n=5 mice/genotype/sex), mice received a single intraperitoneal injection of bromodeoxyuridine (BrdU, 200 mg/kg), a thymidine analog that is incorporated into any cell undergoing DNA synthesis, 2 h before perfusion and were subsequently processed using the same protocol described for morphology studies. All procedures were performed in accordance with the ethical guidelines set forth by the Canadian Council on Animal Care and approved by the Animal Care Committee at the University of British Columbia.

### METHOD DETAILS

#### Microglia isolations, library preparation and sequencing

Brain regions (HC and CTX) were chopped into small pieces and digested with 500-1000 μL digestion buffer (100 μg/mL DNase I, 20 units/sample activated Papain solution, and the inhibitor cocktail) per brain region and digested on ice for 30 minutes. The digested brain solution was further dounced until homogeneous and enriched for microglia using a discontinuous Percoll gradient (20%, 37%, 70%) (Millipore Sigma, GE17-0891-02). The interface between the 37% and 70% layers was collected, and cells were stained with 1:100 P2ry12-APC (Biolegend 848006) and 1:100 Zombie Aqua (biolegend 423101). Live microglia (Zombie aqua-/P2ry12+) were sorted on the BC Cytoflex SRT (Figure S1), and cells were kept on ice until RNA isolation. RNA was collected from sorted cells using the Qiagen RNeasy Plus Micro kit (Qiagen #74034) and the RNA quality was assessed by Bioanalyzer (Agilent 2100 Bioanalyzer, RNA 6000 Pico Kit (5067-1513)). Samples with >5ng of total RNA and RIN of >9 were used for library preparation. (Table S1) Libraries for ribodepleted RNA-sequencing were prepared using Illumina RiboZero Kit and the Illumina Stranded Total RNA prep kit with 13-18 PCR cycles and sequenced at a depth of 40 million pair-end reads using the Illumina HiSeq2000.

#### Histology and Immunohistochemistry

Brains from morphology and validation studies were sliced into 40 µm coronal sections using the Leica SM2000R microtome (Richmond Hill, ON, Canada). The prefrontal cortex and cerebellum were blocked, and hippocampal slices were collected in a series of 8 throughout the extent of the rostral-caudal HC. Slices were stored at -20℃ in 1.5 mL tubes containing antifreeze solution (ethylene glycol, glycerol, and 0.1 M PBS) to further prevent tissue damage due to freezing.

#### Ionized calcium-binding adaptor molecule (Iba1) Expression

To analyze microglia morphology, brain sections from morphology studies were stained for immunofluorescence with a commonly used microglia marker, Ionized calcium-binding adaptor molecule 1 (Iba1) a microglia marker.^69^ Brain sections were transferred into 12-well plates and washed three times for 10 minutes with 0.1 M freshly made phosphate buffered saline (PBS; Sigma-Aldrich, Oakville, ON, Canada) followed by three 10 minute washes in 0.4 % PBS-T (0.1M PBS + 0.4% Triton-X). Tissues were transferred to a blocking solution composed of 5% Normal donkey serum and 0.4% Triton-X in 0.1M PBS for 1 hour. Sections were then incubated in a primary antibody solution containing rabbit anti-Iba1 (1:1000) in 0.4% Triton-X and 5% NDS in 0.1 M PBS overnight at room temperature. Sections were washed three times for 10 minutes in 0.4% PBS-T and then incubated in a secondary antibody solution containing 1:1000 Donkey anti-rabbit Alexa Fluor 488 in 0.4% Triton-X and 5% NDS in 0.1 M PBS for 2 hours at room temperature.

Sections were washed three times for 10 minutes in PBS, adding DAPI at 1:10,000 during the second wash. Sections were mounted onto Superfrost Plus microscope slides and cover-slipped with made PVA DABCO.

#### BrdU/Iba1 double labelling for validation studies

To analyze the proliferation of microglia, brain sections from validation studies were double-labeled with BrdU and Iba1. Briefly, brain sections were washed three times in 0.1M TBS and then blocked for 1.5 hours in a solution composed of 5% Normal Donkey Serum (NDS; MilliporeSigma, Burlington, MA, USA) and 0.3% Triton-X (Sigma-Aldrich) in 0.1M TBS. Sections were incubated overnight at 4°C in a primary antibody solution containing rabbit anti-Iba1 (1:1000) in 0.3% Triton-X and 5% NDS in 0.1 M TBS. Next, sections were washed three times with 0.1M TBS and incubated overnight at 4°C in a secondary antibody solution containing 1:1000 donkey anti-rabbit Alexa Fluor 488 in 0.1M TBS. Sections were then washed three times with TBS, fixed in 4% paraformaldehyde for 10 min, and rinsed twice in 0.9% NaCL for 10 min. DNA was denatured by incubation in 2N HCl for 30 min at 37°C, followed by three washes in TBS. Sections were blocked again for 1.5 hours in the same blocking solution and incubated overnight at 4°C in a BrdU primary antibody solution consisting of 1:1000 anti-BrdU, 0.3% Triton-X and 5% NDS in 0.1 M TBS. After three washes in TBS, sections were incubated overnight at 4°C in a secondary antibody solution containing donkey anti-rat Alexa Fluor 594 (1:500) in 0.1M TBS. Finally, sections were rinsed three times in TBS, mounted onto microscope slides, and cover-slipped using made PVA-DABCO.

#### Imaging analysis of microglia morphology

Brains were imaged at 20x magnification using a ZEISS Axioscan 7 microscope slide scanner. Z-stack images were acquired with a 0.5 µm step size, and Extended Depth of Focus (EDF) images were created using the variance setting within the imaging software (ZEISS ZEN 3.9). Outputs were saved as .czi files and exported as TIFFS. Regions of interest (ROIs) including the HC and CTX were aligned to the Allen Brain Atlas (mouse.brain-map.org) using the ImageJ macro FASTMAP.^70^

These segmented regions were used as input for downstream morphological analysis in MicrogliaMorphology, which uses a Image-J plugins Skeletonize (2D/3D) and AnalyzeSkeleton (2D/3D)^71^ to collect 27 metrics of morphology per microglia cell as detailed in Kim et al., 2024.^46^ The parameters used for MicrogliaMorphology included: auto local thresholding using niblack method and radius 100 with an area filter of 59.6316-775.0912 μm^2^.

#### Imaging analysis of BrdU/Iba1 immunostaining for validation studies

BrdU/Iba1 co-expressing cells were quantified in the dorsal and ventral HC by an experimenter blind to the group assignment (n=2 section/region/mouse; n=5 mice/group) using an Olympus Fluoview 4000 confocal microscope. Cells were counted separately in each region because they are associated with different functions: the dorsal HC with spatial learning and memory and the ventral HC with stress and anxiety.^72,73^ To quantify BrdU/Iba1 co-labelled cells, confocal z-stack images were acquired at 20x magnification with a 10.44 µm range and a 0.87 µm step size. Cells co-expressing BrdU and Iba1 were manually counted, then maximum intensity projections were generated in CellSens and exported as TIFF files. The area of the ROIs (dorsal and ventral HC) was measured using FIJI. BrdU/Iba1co-expressing cell density was calculated dividing the total number of cells by area (µm^2^) of the corresponding ROI.

### QUANTIFICATION AND STATISTICAL ANALYSIS

#### Weight Gene Co-expression Network Analysis (WGCNA)

Considering the complexity of our experimental design, we used WGCNA, a network-based approach that offers key advantages over pairwise differential expression analysis by simultaneously considering all genes across all samples, enabling the identification of coordinated co-expression modules that reflect shared biological functions.^74^ Gene counts were first filtered to include only genes with counts ≥15 in at least 3 of the samples, then normalized and transformed using the variance stabilizing transformation (*vst)* function of the DESeq2 package. Batch effects were corrected using ComBat.^75^ Sample quality control identified one HC sample outlier (hAPOEε3 female), which was excluded from downstream analysis. WGCNA was then performed using the top 50% most variable genes, with separate networks constructed for HC and CTX samples due to brain region differences.^74^ Based on the relationship between power and scale independence, a power of 11 was chosen for HC and 10 for CTX to build scale-free topology using signed networks and biweight midcorrelation (bicor). The minimum module size was set to 40, and modules whose correlation coefficients were greater than 0.65 (mergeCutHeight = 0.35) were merged. Each module was summarized by the first principal component of the scaled module expression profile, termed module eigengene (ME). First, MEs were correlated with four binary traits of interest defined by: hAPOEε4 female (Fε4), hAPOEε3 female (Fε3), hAPOEε4 male (Mε4) and hAPOEε3 male (Mε3). Then, each ME was analyzed using a linear model with Sex, Genotype, and their interaction as fixed factors. Model effects were evaluated using Type III ANOVA and posthoc analyses of genotype within sex were conducted using Šídák’s multiple comparisons test. Significance was set at *p* < 0.05 and effect sizes were reported as partial η². Modules were annotated for Gene Ontology terms (GO) and Kyoto Encyclopedia of Genes and Genomes (KEGG) pathways using R package clusterProfiler (Xu et al., 2024), with *p*-values adjusted by Benjamini-Hochberg (BH) method and significance set at *p* < 0.05. Finally, intramodular hub genes, which are genes with the highest connectivity to other genes within a given module, were selected based on intramodular connectivity (kWithin) and p values of Module membership (MM). Boxplots were used to visualize the distribution by group and gene-gene connections among top hub genes were visualized using Cytoscape version 3.10.3.

#### MGEnrichment analysis

Gene list enrichment analysis was performed in all significant modules identified using MGEnrichment,^29^ with all genes interrogated in the experiment as background. Significant enrichment against mouse microglia gene lists in MGEnrichment were tested by one-tailed Fisher’s exact test and False Discovery Rate (FDR)-corrected to reach significant at an adjusted *p* < 0.05. Target % values for significant gene list enrichment were plotted using ggplot in R.

#### Differential gene expression analysis

Differentially expressed genes (DEGs) were identified in each brain region (HC and CTX) using R package DESeq2 on filtered counts (genes with ≥15 counts in at least 3 samples).^76^ DESeq2 model counts using negative binomial distribution, with variance and mean linked by parametric regression. The model design included the interaction between sex and APOE genotype, as well as CollectionDate to correct for batch effect. Contrasts of interest were extracted to identify DEGs for female APOE4 vs. APOE3 and male APOE4 vs. APOE3 comparisons. The sex by genotype interaction term was also extracted. For each contrast, the base mean expression, the log2 fold change, unadjusted p-value and q-values (FDR-corrected p value) were calculated, and genes with q-value < 0.05 were considered significant. Heatmaps were used to visualize expression patterns of DEGs, and UpSet plots were used to visualize the overlap between DEGs identified in each contrast of interest and the gene modules detected by WGCNA.

#### Microglia morphology analysis

MicrogliaMorphologyR (R package) was used to analyze and visualize our dataset, as described in Kim et al., 2024,^46^ in order to characterize microglia morphological states and gain insight into their relevance in experimental models. Briefly, dimensionality reduction was conducted using principal component analysis followed by k-mean clustering on the first three principal components to define four morphological classes within our dataset (ramified, hypertrophic, ameboid and rod-like). The optimal clustering parameter was identified using exploratory data analysis methods including the within sum of squares and silhouette methods.

Cluster identities were assigned by examining the relationship between each cluster and the 27 morphological features and by using the ColorByCluster features in MicrogliaMorphology to visually examine the different cluster morphologies.

To assess how the percentage of cells in each morphology cluster changed with APOE genotype in a sex-dependent manner within each brain region, a generalized linear mixed model was fitted using the glmmTMB R package,^77^ to measure cluster percentage change as a factor of Cluster identity, APOE genotype and sex interaction with MouseID as a repeated measure (percentage ∼ Cluster*Genotype*Sex + (1|MouseID)). The analysis was performed using the MicrogliamorphologyR, which calls the glmmTMB R package.^77^ Model effects were evaluated using three-way Analysis of Deviance (Type II Wald chisquare test) test and pairwise test between APOE Genotype groups were corrected for multiple comparisons across Clusters and Sex (∼Genotype|Cluster|Sex) using the Šídák’s method and adjusted *p* < 0.05 were considered statistically significant.

#### Statistical analyses

Data from microglia proliferation were expressed as mean ± standard error of the mean (SEM) and analysed with STATISTICA (StatSoft). Data were tested for normality using the Shapiro-Wilk test and for homogeneity of variance using the Levene test. BrdU/Iba1 co-expressing cells were each analyzed using repeated-measures ANOVA, with genotype (hAPOEε3 and hAPOEε4) and sex (male and female) as between-subject variables and with hippocampal region (dorsal, ventral) as the within-subject variable. *A priori* comparisons were subjected to Bonferroni corrections. Significance was set to α = 0.05 and effect sizes are given with Cohen’s *d* or partial η^2^.

