## Supplement Figures and Table captions for "Sex and APOE Genotype Differentially Shape Microglial Transcriptomic Profiles Across the Hippocampus and Cortex"

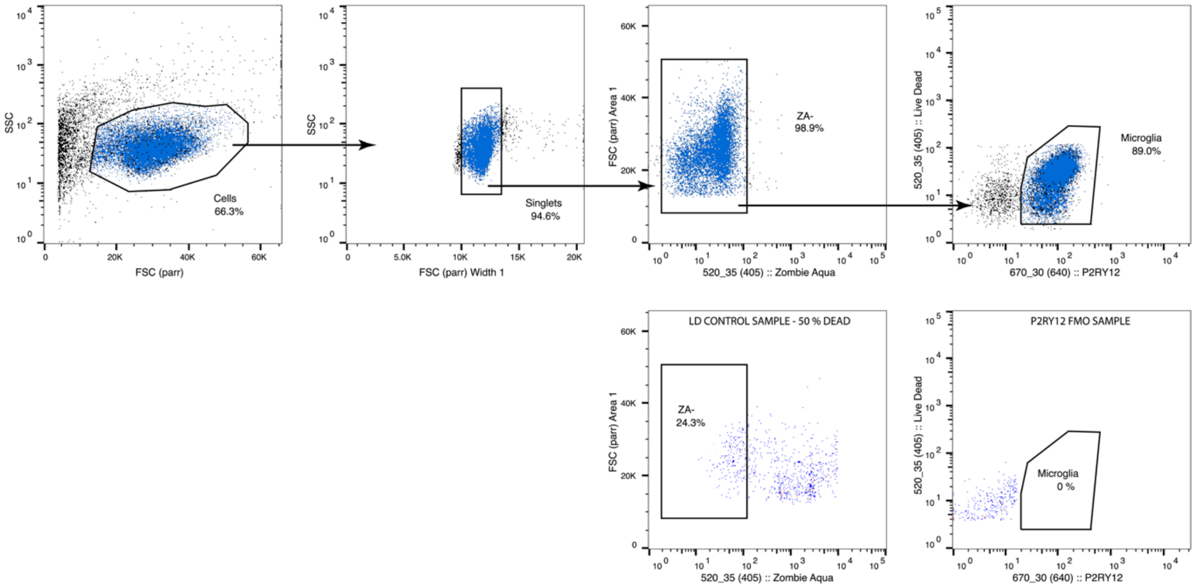


**Figure S1**. **Gating Strategy for Microglia Isolation.** Microglia were first identified by shape as singlet cells (SSC x FSC; SSC vs FSC-Width), then as Zombia-Aqua (ZA) negative based on V520 and P2RY12-APC positive on R670. ZA was gated on a killed control (24.3% live) and P2RY12 was gated on an FMO. Cells were sorted for purity on a BD Influx II machine with sample pressure of ~18psi. Average efficiency for each sort was ~95%.

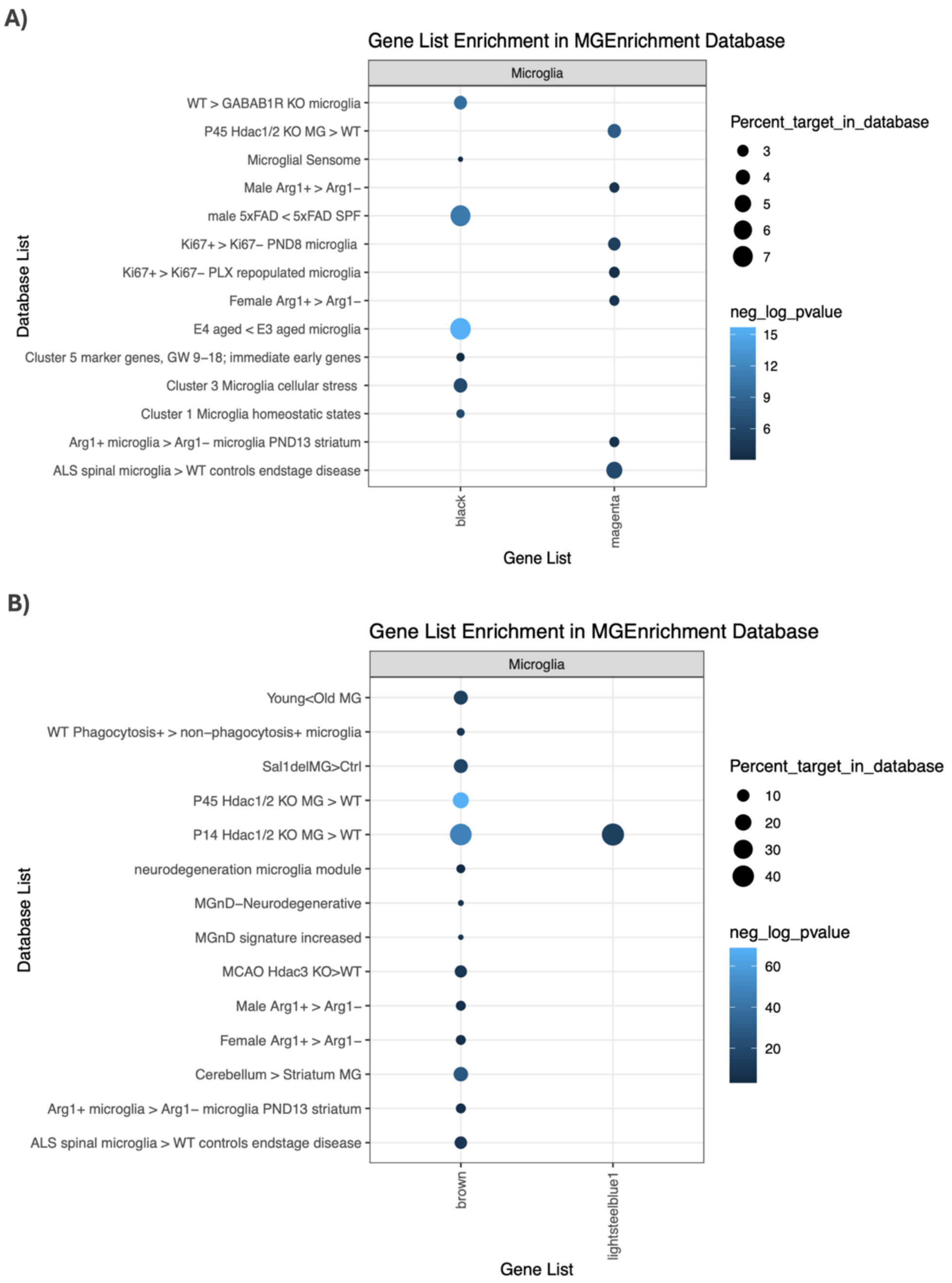


**Figure S2. Black, magenta, brown and lightsteelblue1 WGCNA modules are enriched for microglia-relevant signatures.** (A,B) Significant enrichments against mouse microglial gene lists in MGEnrichment in (A) the hippocampus and (B) the cortex. Blue circles denote a significant enrichment in the percent of genes within each module for a given microglia gene list compared to that of the background of all genes detected in the RNA-Seq experiment. Color intensity represents the −log(FDR-adjusted p-value), with lighter blue indicating greater statistical significance.

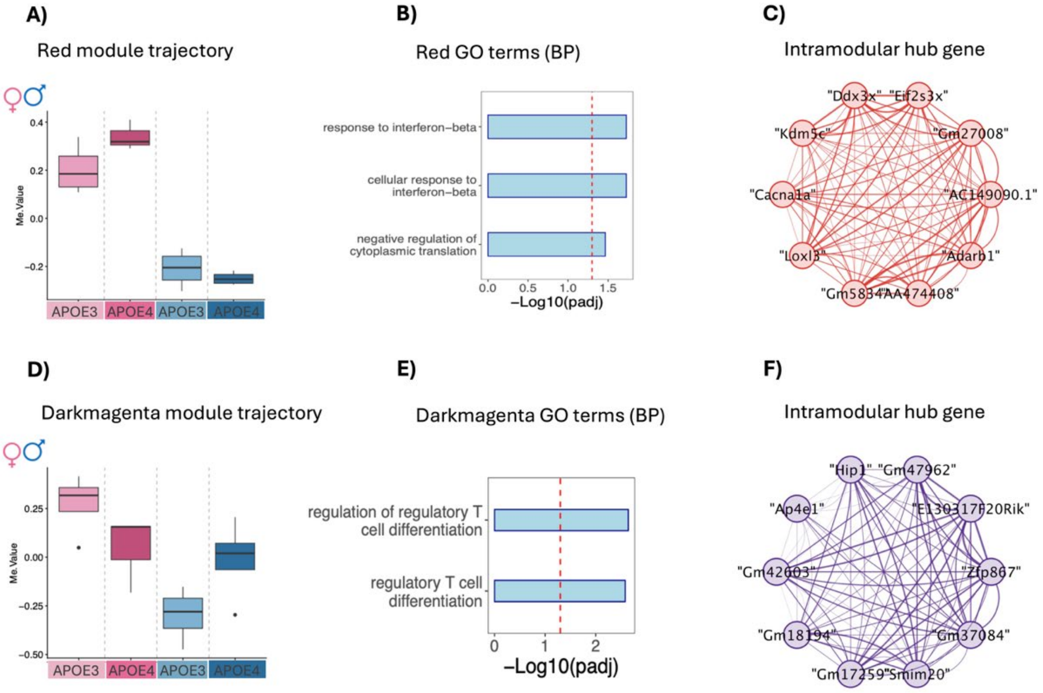


**Figure S3. The interferon-enriched red module and the T cell differentiation-enriched darkmagenta module both showed a significant sex-by-genotype interaction. (A,D)** Module eigengene boxplot in the **(A)** red and **(D)** darkmagenta modules across APOE genotype and sex. **(B, E)** The top 5 Gene Ontology (GO) enriched Biological Processes (BP) in **(B)** red and **(E)** darkmagenta modules. The dotted line indicates the threshold for BH-corrected p-values < 0.05. **(C, F)** Network plot with the top 10 intramodular hub genes in the **(C)** red and **(F)** darkmagenta modules.

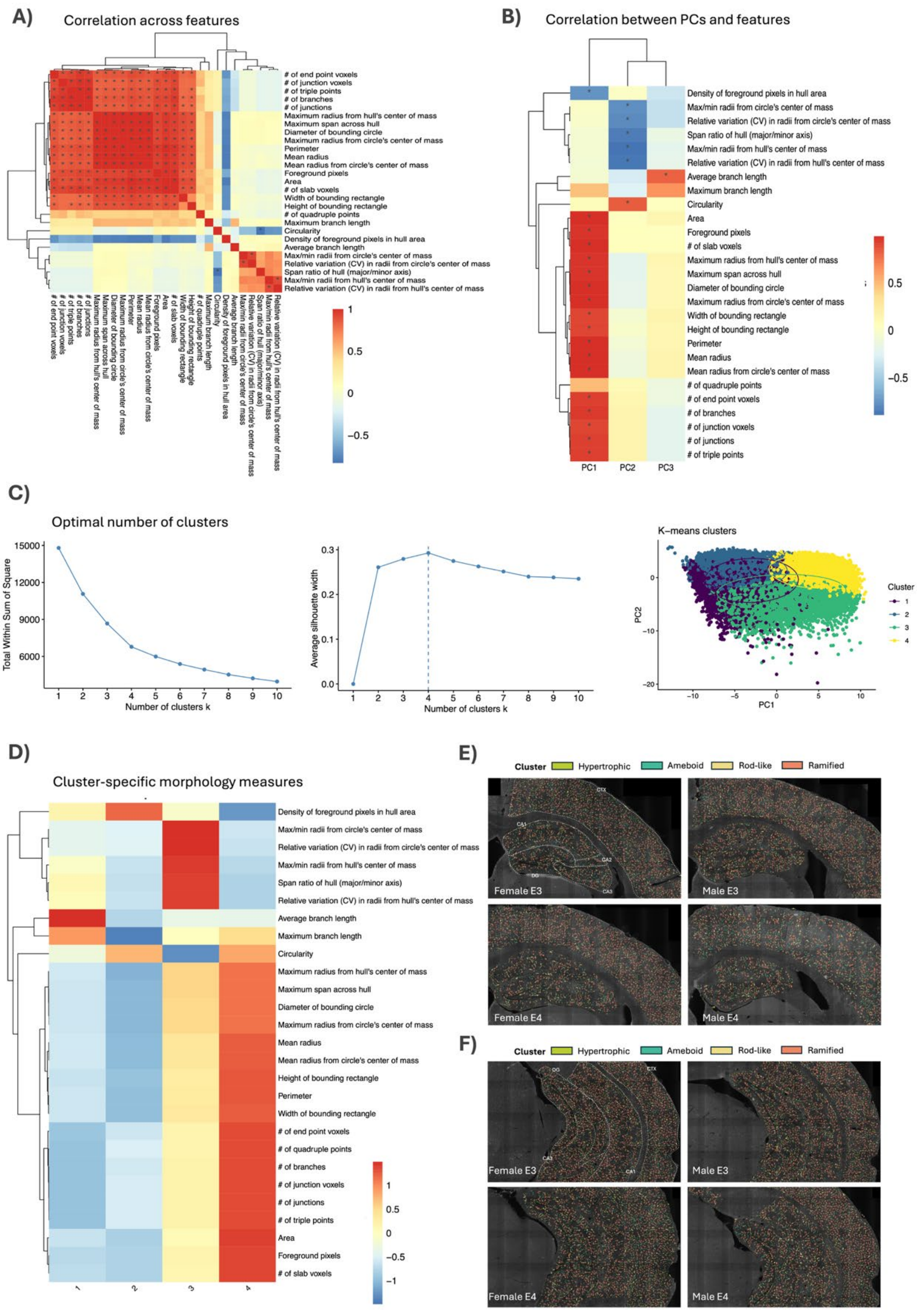


**Figure S4.** Analysis of microglia morphology state using MicrogliaMorphology and MicrogliaMorphologyR. **(A)** Spearman's correlation matrix of 27 features measured by MicrogliaMorphology (*$p < 0.05$). **(B)** Spearman's correlation of morphology measures to first 3 principal components after dimensionality reduction(*$p < 0.05$). **(C)** Optimal k-mean clusters determined using sum of squares and silhouette. Cluster classes displayed in PCs 1–2 space. **(D)** Average values for all 27 morphology features, centered and scaled across clusters. **(E-F)** Individual cells spatially registered back to original images and visually annotated by morphological class using ColorByCluster feature in **(E)** dorsal hippocampus and **(F)** ventral hippocampus.

| Sample # | Brain Region | Sex | Genotype | RIN | Total RNA (ng) |
|---|---|---|---|---|---|
| 1 | HPC | male | hAPOE3 | 9.5 | 15.384 |
| 2 | HPC | male | hAPOE3 | 9.7 | 15.216 |
| 3 | HPC | female | hAPOE3 | 9.2 | 14.028 |
| 4 | HPC | female | hAPOE4 | 9.3 | 14.892 |
| 5 | HPC | female | hAPOE4 | 9.6 | 11.184 |
| 6 | HPC | male | hAPOE4 | 9.5 | 13.776 |
| 7 | HPC | male | hAPOE4 | 9.9 | 9.108 |
| 8 | HPC | female | hAPOE3 | 9.4 | 13.476 |
| 9 | HPC | female | hAPOE4 | 9.9 | 9.708 |
| 10 | HPC | female | hAPOE4 | 9.8 | 11.016 |
| 11 | HPC | male | hAPOE3 | 9.7 | 7.536 |
| 12 | HPC | male | hAPOE3 | 10 | 10.044 |
| 14 | HPC | female | hAPOE3 | 9.7 | 6.312 |
| 15 | HPC | male | hAPOE4 | 10 | 14.28 |
| 16 | HPC | male | hAPOE4 | 10 | 6.108 |
| 1 | CTX | male | hAPOE3 | 9.4 | 31.776 |
| 2 | CTX | male | hAPOE3 | 9.3 | 47.568 |
| 3 | CTX | female | hAPOE3 | 9.5 | 16.884 |
| 4 | CTX | female | hAPOE4 | 9.8 | 16.092 |
| 6 | CTX | male | hAPOE4 | 9.8 | 13.776 |
| 7 | CTX | male | hAPOE4 | 9.7 | 6.576 |
| 8 | CTX | female | hAPOE3 | 9.5 | 12.972 |
| 9 | CTX | female | hAPOE4 | 9.9 | 8.22 |
| 10 | CTX | female | hAPOE4 | 9.8 | 14.772 |
| 11 | CTX | male | hAPOE3 | 9.6 | 15.852 |
| 12 | CTX | male | hAPOE3 | 9.8 | 10.68 |
| 13 | CTX | female | hAPOE3 | 9.6 | 12.168 |
| 14 | CTX | female | hAPOE3 | 9 | 6.12 |
| 15 | CTX | male | hAPOE4 | 9.7 | 15.9 |
| 16 | CTX | male | hAPOE4 | 9.6 | 13.536 |

**Table S1.** RNA Integrity Number (RIN) and total input RNA (ng) for RNA-seq samples

**Table S2.** WGCNA module information from HC, Pearson's correlation between module eigengene (ME) and groups of interests, linear model results, posthoc analysis results, Gene ontology (GO) terms, Kyoto Encyclopedia of Genes and Genomes (KEGG) and microglial gene set enrichments (MEnrichment) related to Figure 1.

**Table S3.** Differentially expression analysis results from hippocampal microglia related to Figure 3.

**Table S4.** WGCNA module information from CTX, Pearson's correlation between module eigengene (ME) and groups of interests, linear model results, posthoc analysis results, Gene ontology (GO) terms, Kyoto Encyclopedia of Genes and Genomes (KEGG) and microglial gene set enrichments (MEnrichment) related to Figure 4.

**Table S5.** Differentially expression analysis results from cortical microglia related to Figure 5.

**Table S6.** Type II Wald chisquare test results and Šídák's-corrected posthocs from MicrogliaMorphology cluster analysis related to Figure 6.
